# AURKB/Ipl1 restarts replication forks to recover from replication stress

**DOI:** 10.1101/2021.05.27.446025

**Authors:** Erin Moran, Katherine Pfister, Kizhakke Mattada Sathyan, Brianne Vignero, Daniel J. Burke, P. Todd Stukenberg

**Affiliations:** Department of Biochemistry and Molecular Genetics, University of Virginia, School of Medicine, Charlottesville, VA, 22908; Department of Biological Sciences, North Carolina State University, Raleigh, NC, 27695

## Abstract

Aurora kinase B (AURKB in human, Ipl1 in *S. cerevisiae*) is a master regulator of mitosis and its dysregulation has been implicated in chromosome instability. AURKB accumulates in the nucleus in S-phase and is regulated by CHK1 but has not been implicated in in the DNA Damage Response (DDR). Here we show that AURKB has a conserved role to recover from replication stress and restart replication forks. Active AURKB is localized to replication forks after a prolonged arrest. CHK1 phosphorylation of AURKB induces activating phosphorylation of PLK1 and both Aurora and Plk1 are required to deactivate the DDR. Clinical trials with AURKB inhibitors are designed to target established roles for AURKB in mitosis. Our data suggest combinations of AURKB inhibitors and DNA damaging agents could be of therapeutic importance.

## Introduction

Genomic regions with a paucity of replication origins, regions replicated from unidirectional forks, and regions with convergent replication and transcription are at risk for replication fork “pausing” and rely on fork-restart mechanisms in an unperturbed S-phase^1,2^. Replication forks also stall in response to DNA damage and there are mechanisms to stabilize replisomes and to restart replication from stalled forks. Factors promoting the fork stability and restart are part of the DNA Damage Response (DDR) that includes both the intra-S checkpoint that responds to stalled replication forks and the DNA damage checkpoint that responds to double stranded breaks, requiring homologous recombination to restart stalled forks.

The essential function of the DDR during S-phase is to prevent replication fork collapse and promote replication re-start ^3^. Replication stress, or replication fork pausing, is induced experimentally by depleting ribonucleotides (with hydroxyurea (HU)) or exposing cells to agents that modify the DNA such as ultraviolet (UV) light DNA modifying chemicals such as the alkylating agent methyl methanesulfonate (MMS). UV, MMS and other DNA-modifying chemicals generate physical barrier that induces fork pausing^1^. ssDNA at stalled replication forks is coated with RPA that forms a signaling platform recruiting numerous proteins to activate ATR (Ataxia-Telangectasia and Rad3 Related) kinase in humans or Mec1 kinase homolog in yeast^4^. ATR/Mec1 phosphorylates several proteins at stalled replication forks, represses firing of late origins and activates the effector kinases especially CHK1/Rad53, which promotes cell cycle arrest, regulates dNTP and histones levels to facilitate fork restart and recruits cryptic origins to complete DNA synthesis.

Aurora B kinase (AURKB in human and Ipl1 in yeast) is a master regulator of multiple events in mitosis, and its overexpression leads to chromosome instability in many tumors ^5–7^. AURKB exists in a four subunuit complex know as the Chromosome Passenger Complex (CPC), which also contains the subunits INCENP, Survivin, and Borealin^5^. Suppressing AURKB with chemical inhibitors impairs most CPC functions. However, the non-enzymatic partners of the CPC are essential for proper targeting and timely activation of the kinase ^5,8^. Two histone marks are required to recruit CPC to the inner centromere in mitosis: Survivin directly binds histone H3 after it is phosphorylated on Thr-3 by Haspin kinase^9,10^, while Borealin interacts with Sgo1, which recognizes histone H2A phosphorylation at Thr-120 by Bub1 kinase ^11,12^. AURKB can auto-activate by phosphorylating T-loops in *trans*, which is more efficient than activation in *cis*, suggesting that concentration of the CPC is sufficient to initiate a local signal. However, there is also input from the DDR kinase CHK1 which phosphorylates AURKB on S331 and has a poorly understood role in regulating AURKB activity^13^.

Yeast have a single Aurora kinase, Ipl1, while vertebrates have at least two referred to as Aurora A and Aurora B. Aurora A has been shown to regulate the entry of mitosis where it inactivates the DDR by activating Plk1^14^. Aurora kinases have many substrates in mitosis but a key form of signal transduction is that they can activate another important mitotic regulator Polo Like Kinase 1 (PLK1)/Cdc5 in yeast. Yeast Cdc5 regulates the DDR after prolonged G2 arrest induced by irreparable DNA double-strand breaks and Plk1 can similarly override the DDR in response to replication stress in vertebrates ^15–18^. In human cells, PLK1 has also been shown to phosphorylate Rad51 to facilitate homologous recombination in response to double strand breaks and to phosphorylate MRE1 to inactivate the MRN complex to limit its activity^19,20^.

Here we show that the Aurora B kinase has an evolutionarily conserved role during S phase and is required for cells to recover from replication stress. Active AURKB localizes to stalled replication forks where its activity is required to inactivate the stress response and allow replication forks to restart after the stress is removed. AURKB activity in S-phase requires upstream activation by CHK1 and it is required to activate PLK1 during fork restart. Overall our data establish that Aurora B kinase has important functions in S-phase in the DDR.

## Results

### Cells lacking Aurora kinase activity are sensitive to DNA damaging agents

We determined if there was a role for Ipl1 in the DDR in yeast by growing temperature-sensitive *ipl1-321* cells at the semi-permissive temperature to limit them for Ipl1 activity in the presence or absence of sublethal amounts of hydroxyurea (HU) and methyl-methanesulfonate (MMS). We also treated the cells briefly with ultraviolet light (UV) light before incubating them at the semi-permissive temperature. The *ipl1-321* cells are compromised for growth on YPD medium at 32 degrees showing they are limited for Ipl1 activity compared to the wild type and cells lacking Mec1, the yeast homolog of ATR (Ataxia-Telangectasia and Rad3 Related) kinase in humans. Cells with limited Ipl1 activity are sensitive to UV and to chronic growth in the presence of MMS and HU (Figure 1A). All three treatments have in common that they impose replication stress by blocking replication fork progression and activate the DDR. Yeast cells were also sensitive to brief acute treatments of MMS and viability dropped one hundred-fold after 90 minutes of 0.033% MMS treatment (Supplemental Figure 1A).

**Figure 1.**
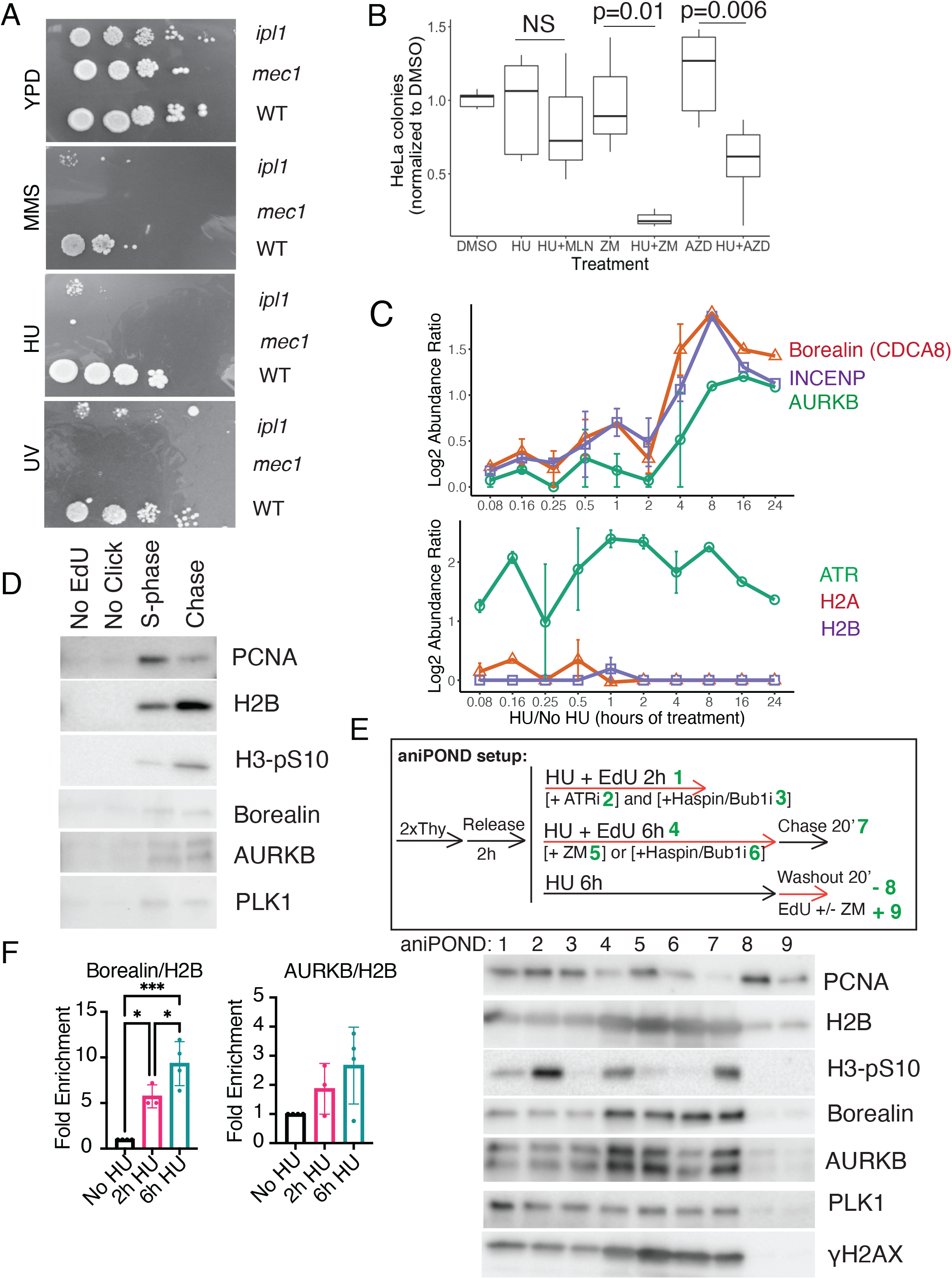
Aurora B is required for recovery from replication stress. AURKB has a role in the response to DNA damage. **A**. Yeast cells limited for Ipl1 are sensitive to DNA damage. Ten-fold serial dilution of cells spotted onto YPD plates and grown continually with 0.01%MMS, 50mM HU, or after 50 uJ/cm^2^ of ultraviolet light. WT (W303), *ipl1* (4145-1-3) and *mec1* (JRY7274) **B**. Replication stressed HeLa cells are sensitive to two distinct AURKB inhibitors but not an AURKA inhibitor. HeLa cells were treated with the indicated drugs (100nM AZD1152 (AZD), ZM-0.4uM ZM447439, 50nM MLN8237 (Alisertib, MLN), 2mM hydroxyurea (HU)) for 6 hours and ∼350 cells were plated in 10cm plates and grown for 15 days. The number of colonies formed were counted after staining with crystal violet. P-values calculated by Anova with Tukey post-test. **C**. Proteins at stalled replication forks. Quantitative iPond SILAC analysis of the indicated proteins Log_2_ Abundance Ratio of HU treated cells over time compared with a non-HU treated control (Data mined from Dungrawala *et al*., 2015^21^). **D**. aniPOND of HeLa cells synchronized into S phase. Cells were pulsed with EdU for 10 minutes, and then chased with 10 μM Thymidine for 20 minutes. Negative controls for the EdU labeling (No EdU) and the conjugation to biotin (No Click) were also included. Samples were subjected to western blotting of the respective proteins after normalization to the total amount of input protein. **E**. aniPOND of HeLa cells synchronized into S phase and incubated with 3 mM HU to induce replication stress. Top, schematic of experiment, with red arrows indicating where EdU is incorporated in the various samples. Bottom, samples are subjected to western blotting of the respective proteins after normalization to the total amount of input protein in each sample. **F**. Quantification of CPC proteins from aniPOND samples, normalized to H2B. P-values calculated by one-way ANOVA with Tukey post-test with multiple comparison correction.

We determined if human cells, limited for AURKB activity, were also sensitive to replication stress. HeLa cells treated with HU for up to eight hours retain high viability but lose viability when grown in the presence of HU and the AURKB inhibitor ZM44739 (ZM) for longer than 4 hours (Figure 1B). The sensitivity of HeLa cells to HU was seen with two different AURKB inhibitors (ZM and AZD1152-HQPA(AZD)) that have distinct chemical structures (Figure 1B). This sensitivity was not seen in cells treated with HU and the AURKA inhibitor MLN8237 (MLN) (Figure 1B) and sensitivity appeared after 6 hours of HU arrest (Supplemental Figure 1B). We confirmed that the concentrations of ZM and AZD used throughout these experiments inhibit AURKB activity but not AURKA activity by immunoblots using a Pan-Aurora kinase T-loop antibody (Supplemental Figure 1E).

### The chromosome passenger complex (CPC) localizes to stalled replication forks

A recent study utilized iPOND (isolation of proteins on nascent DNA) combined with SILAC (stable isotope labeling of amino acids in cell culture)-mass spectrometry (MS) to analyze replisomes at stalled replication forks over time^21^. Human embryonic kidney (HEK293T) cells were treated with hydroxyurea and replication forks were purified using iPOND. This study provides a comprehensive description of the active and stalled replication fork-associated proteomes. We mined the quantitative proteomic data set and found that CPC is recruited to stalled replication forks. The CPC proteins AURKB, Borealin (CDCA8), and INCENP accumulated at the stalled forks and their proportion increased between four and eight hours of hydroxyurea treatment (Figure 1C). We used a subset of the quantitative proteomic data to get those proteins that were enriched at the later time points and applied unsupervised hierarchical clustering to determine the similarities in the accumulation of proteins at stalled forks (a subset of the data is shown in Supplemental Figure 1C). Borealin (CDCA8) and INCENP cluster immediately beside each other and are similar to proteins like BLM, WRN and SMARCAL1 that are required to stabilize replication forks. AURKB and PLK1 also cluster near each other and this suggests that there is a role for the mitotic kinases in the response to replication stress.

We confirmed that the CPC is recruited to stalled replication forks, but is not enriched at nascent forks by aniPOND (Figure 1D-F, Supplemental Figure 1D^22^). S Phase HeLa cells were labeled for 10 minutes with EdU and nascent replication forks were isolated from chromatin and associated proteins were detected by immunoblotting. Nascent replication forks contain Proliferating Cell Nuclear Antigen (PCNA). We identified AURKB, Borealin, and PLK1 present at nascent replicating forks, but not present within the negative control unlabeled (No EdU) or non-conjugated to biotin (No Click) samples. We also found serine 10-phosphorylated histone H3 (H3pS10), a known substrate of active AURKB, present at nascent replicating forks. However, these signals were still present after a 20-minute chase of the EdU signal away from the nascent fork (Chase), unlike PCNA, indicating that the CPC is present at replicating chromatin but not enriched at nascent replication forks. We also localized the CPC to replicating chromatin during and after replication stress (Figure 1E). We measured higher levels of both Borealin and AURKB as well as AURKB activity on S10 of H3 on forks arrested for 6 hours in HU than 2 hours (Figure 1E lanes 1 and 4, Figure 1F) consistent with the aforementioned iPOND data^21^. Note the H3-pS10 activity at stressed forks depends on AURKB activity as it was lost after the addition of the AURKB inhibitor ZM447439 (ZM) (Figure 1E, lanes 4, 5). These pools of CPC remained on the stressed DNA after removal of the HU to restart replication forks suggesting that the CPC remains at the chromatin that was stressed (Figure 1E, lane 7). The increased recruitment of activated CPC to forks arrested for 6 hours did not depend on AURKB activity although it did depend on the Haspin and BUB1 histone kinases that localize the CPC to inner centromeres during mitosis (Figure1 E lanes 4,5,6). Surprisingly, if we added an ATR inhibitor (VE822) to forks arrested in HU for two hours to induce fork collapse we detected an increase in H3-pS10 activity (Figure 1E lanes 1 and 2)). These data suggest that AURKB responds to collapsed forks. Cells were washed out of HU into EdU to determine if the CPC moved with restarted replication forks after being stressed and little CPC was found on restarted forks. Treating restarting forks with an AURKB inhibitor resulted in less PCNA recruitment suggesting a role for AURKB in replication restart (Figure1E, lanes 8,9). We validated the effects of Haspin, BUB1, and AURKB inhibitors on documented histone targets H3-pT3, H2A-pT120, and H3-pS10 respectively in Supplemental Figure 3C.

### Aurora kinases are required restart stalled replication forks

We used an “analog-sensitive” *ipl1* mutant (*ipl1-as*) to determine how MMS affects DNA synthesis in yeast cells in the absence of Ipl1^23^. The protein kinase activity of Ipl1 in cells with the *ipl1-as* mutation is inhibited by the addition of the ATP analogue 1-naphthyl-pyrazolo[3,4-d]pyrimidine (NAPP). We arrested *ipl1-as* cells with the mating pheromone α-factor and sampled cells for flow cytometry at twenty-minute intervals in the presence and absence of MMS and NAPP. Untreated and NAPP-treated cells replicated their DNA with kinetics that are similar to wild type cells (Figure 2A). Cells treated with 0.033% MMS progressed slowly through S phase and accumulate with a G2/M content of DNA as expected for cells with an intact DDR. The *ipl1-as* cells grown in the presence of both MMS and NAPP retained a G1-like content of DNA. Thus, the slow movement of DNA replication in MMS-treated cells is dependent upon Ipl1. The effect was not specific to *ipl1-as* cells and NAPP as we obtained similar results using temperature-sensitive *ipl1-321* cells (Supplemental Figure 2A). Importantly, replication in cells lacking Ipl1 appears normal in the absence of replication stress (Figure 2A, Supplemental Figure 2A). The delay in DNA replication in response to MMS in cells lacking Ipl1 is dependent on Mec1 (Supplemental Figure 2B) but independent of the spindle assembly checkpoint (*mad2::KanMx4*) and kinetochores (*ndc10-1*), both of which are regulated by AURKB (Supplemental Figure 2 C,D). We analyzed the ip*l1-as* cells for budding, which is an indication of B-type cyclin activity that is required for both budding and DNA replication. The *ipl1-as* cells treated with NAPP were budded suggesting that the cells with a G1-like content of DNA had cyclin B activity (Supplemental Figure 2E). We used bromo-deoxyuridine (BrdU) labeling to determine if temperature-sensitive *ipl1-321* cells lacking Ipl1 activity initiate DNA replication. We synchronized *ipl1-321* cells in G1 at the permissive temperature with α-factor, sampled cells after releasing them to the restrictive temperature in the presence and absence of MMS. Chromosomal DNA was extracted and separated by contour-clamped homogeneous electric field (CHEF) electrophoresis which separates intact chromosomes. DNA was separated, blotted to membranes and probed for BrdU incorporation with anti-BrdU antibody. There was no BrdU incorporation in α-factor-arrested cells as expected (Figure 2B). Untreated cells lacking Ipl1 incorporate BrdU and intact chromosomes become visible as cells complete replication. Incompletely replicated chromosomes remain in the wells of the CHEF gels. MMS-treated *ipl1-321* cells incorporated BrdU but intact chromosomes were not observed. We confirmed that *ipl1-321* cells initiate replication when treated with MMS by immunoprecipitating BrdU followed by qPCR. DNA replication in yeast initiates from well characterized ARS sequences in a temporal pattern such that an ARS can be described as early or late replicating^24^. We measured BrdU incorporation into two early ARS regions by immunoprecipitating newly incorporated BrdU from cells released from α-factor plus MMS for one and six hours^24^. We used quantitative PCR (qPCR) to determine the extent of BrdU incorporation in ARS sequences. Wild type cells treated for one hour with MMS incorporated BrdU in two early ARS sequences and much less into late ARS sequences (Figure 2C). We could detect approximately equal incorporation into early and late ARS sequences by six hours. We measured BrdU incorporation in the two early ARS sequences and much less into late ARS sequences in *ipl1* cells after one hour and the same difference was seen between early and late ARS sequences after six hours. Cells lacking Ipl1 replicate DNA, however their replication is severely impeded in response to replication stress.

**Figure 2.**
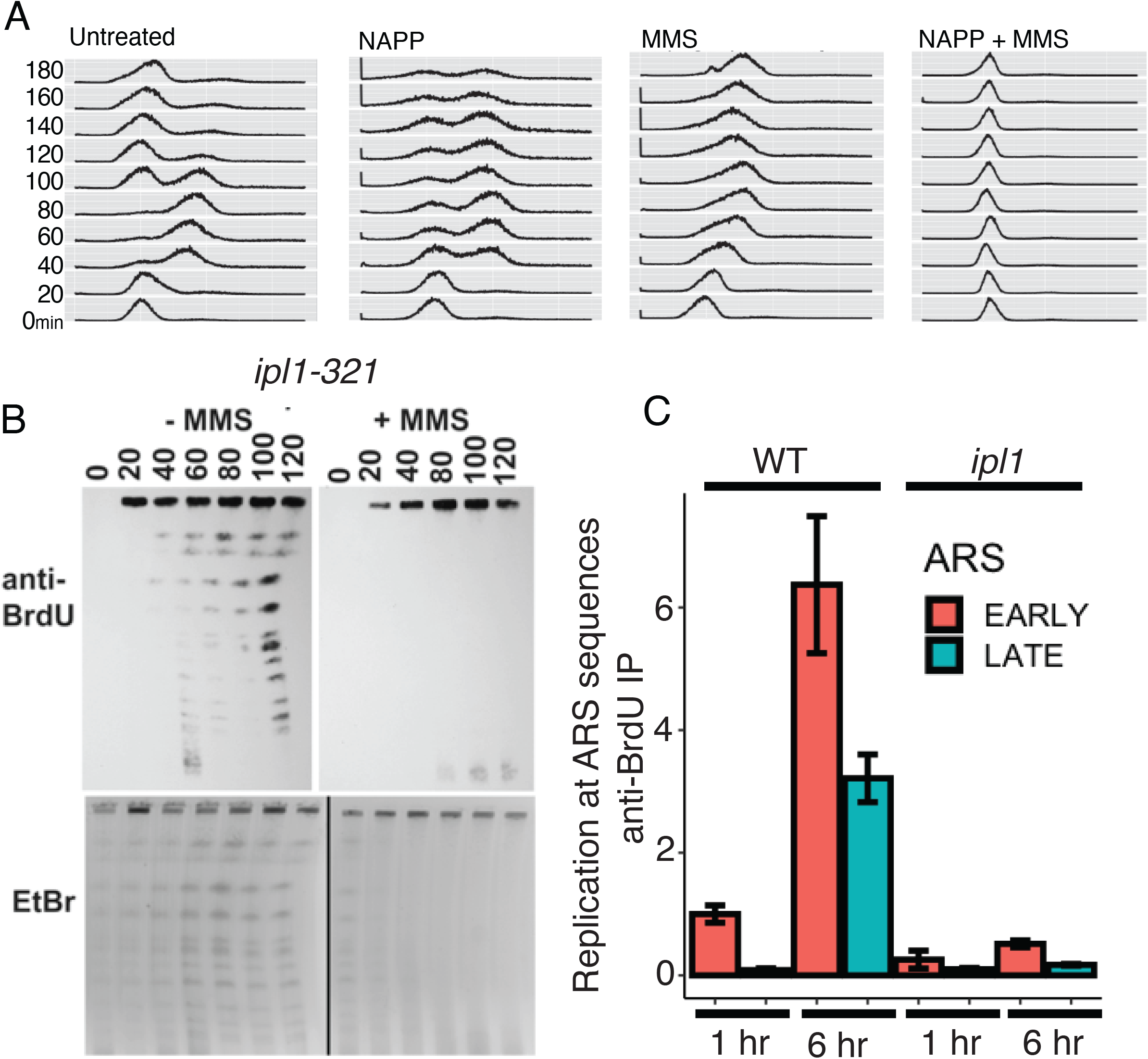
Ipl1/Aurora is required to replicate DNA under replication stress conditions in *S. cerevisiae*. Ipl1 is required to complete replication in the presence of MMS. **A**. Flow cytometry. α-Factor-arrested *ipl1-as* cells (2946-14-2) were released into the cell cycle with the indicated treatment and samples were taken every 20 min for flow cytometry. **B**. Cells lacking Ipl1 enter S-phase but cannot complete chromosome replication. α-Factor-arrested *ipl1-321* cells (2498-3-3) in BrdU were released into the cell cycle for the indicated time and chromosomes were separated by a CHEF gel. DNA was detected with ethidium bromide and BrdU incorporation was determined by immunoblotting. **C**. Origins of replication are activated slowly in Ipl1 cells in the presence of MMS. α-Factor-arrested wild type (CVY43) and *ipl1-321* cells (2498-3-3) were released into the cell cycle in the presence of MMS for 1 and 6 hours. BrdU-containing DNA was recovered by immunoprecipitation and ARS sequences were identified by qPCR. Early ARS sequences were ARS1 and ARS305 and late ARS sequences were ARS440 and ARS522. Data are means and SEM of the amount of DNA recovered for the early and late ARS sequences normalized to the one-hour sample from wild type cells.

We hypothesized that Aurora functions to restart stalled replication forks because it is enriched at stalled forks in human cells eight hours after replication is arrested with HU (Figure 1C). We used flow cytometry to determine if MMS-treated yeast cells lacking Ipl1 activity could efficiently restart replication. Wild type and *ipl1-321* cells were arrested with α-factor and released to the cell cycle in the presence of MMS for 45 minutes and then the MMS was removed (restart). Wild type cells restart from MMS treatment within 60 minutes and the majority of cells completed DNA replication and had a 2C content of DNA (Figure 3A). The *ipl1-321* cells restart from MMS treatment more slowly suggesting that Ipl1 is required for efficient restart from replication stress. This was confirmed by directly measuring total BrdU incorporation during restart (Supplemental Figure 3A) and by qPCR, following immunoprecipitation of BrdU, for early and late ARS sequences (Figure 3B). BrdU was detected at both early and late ARS sequences sixty minutes after restart in wild type cells but BrdU is detected in early ARS sequences but not detected at late ARS sequences in *ipl1* cells.

**Figure 3.**
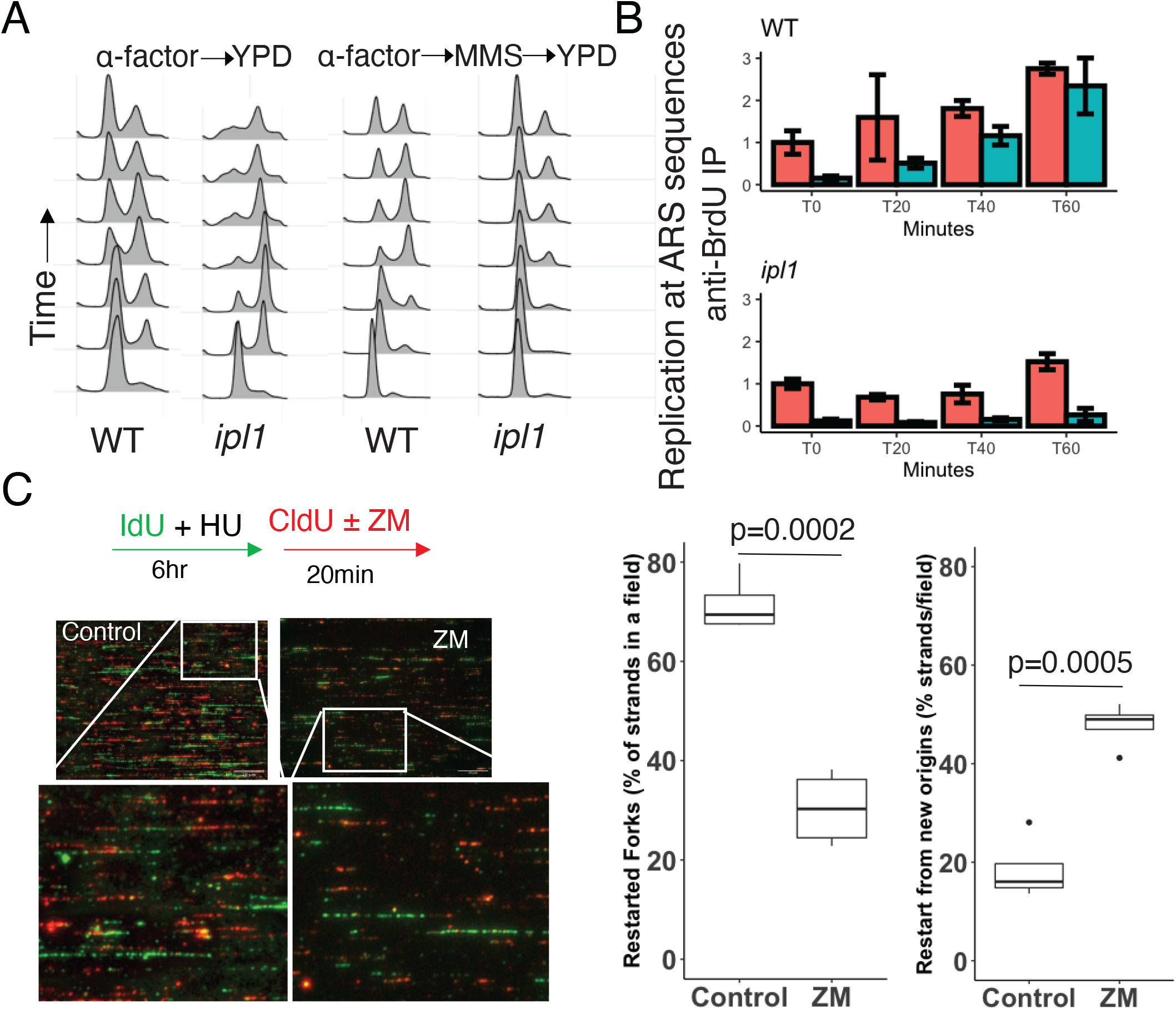
Ipl1/Aurora B is required to restart replication forks after release from replication stress. AURKB/Ipl1 kinases are required for efficient replication restart. **A**. Wild type (CVY43) and *ipl1-321* (2948-3-3) cells were α-factor-arrested and released into the cell cycle for the indicated time at 36 degrees and samples were taken every twenty minutes for flow cytometry. Cells on the right were released in the presence of MMS for thirty minutes and the MMS was removed to assay replication restart. **B**. Late origins of replication are delayed after replication restart in *ipl1* cells. α-Factor-arrested wild type (CVY43) and *ipl1-321* cells (2498-3-3) were released into the cell cycle as in Figure 3A and BrdU-containing DNA was recovered by immunoprecipitation. ARS sequences were identified by qPCR as in Figure 2B. Data are means and SEM of the amount of DNA recovered for the early and late ARS sequences normalized to the one-hour sample. **C**. HeLa cells fail to properly restart replication forks in the presence of AURKB inhibitors. HeLa cells were incubated with IdU nucleoside and HU and then restarted in the presence of CldU in the presence (n=4 fields) and absence of ZM (n=4) and the replicated DNA strands were visualized by DNA combing. Statistics are students t-test.

We determined that AURKB was required for restarting DNA replication during recovery from replication stress in human cells (HeLa) using DNA combing. Cells were synchronized at the beginning of S-phase, released into the cell cycle for one hour and then treated with HU for six hours in the presence of the thymidine analog Iododeoxyuridine (IdU). Cells were then washed out of the HU for twenty minutes in the presence of chlorodeoxyuridine (CldU) in the presence and absence of the AURKB inhibitor ZM. Isolated DNA was stretched on silanized coverslips to separate individual DNA strands (DNA combing) and the DNA synthesized during and after the HU treatment were visualized using specific antibodies (Figure 3C). Replication restart is evident by red stands that are adjacent to green strands. Forks that fail to restart are identified by green strands that lacked or had very short adjacent red strands. The proportion of replicons that restarted were reduced in cells that were treated with AURKB inhibitor ZM (Figure 3C). We found a higher proportion of strands that were only red in ZM treated cells indicative of forks initiated at new origins (Figure 3C). We also observed a smaller proportion of forks restarting in a DNA fiber assay after restarting forks with AZD (Supplemental Figure 3B), demonstrating that two different inhibitors of AURKB affect restart. We performed a restart experiment in human cells and used a western blot to observe γH2AX after restart with AURKB inhibitors in order to address the possibility that we were inducing double strand breaks after restart with these inhibitors. We observed a trend towards lower γH2AX with these inhibitors, which was not significantly different than controls (Supplemental Figure 3C). We conclude that AURKB is required to efficiently restart replication from a stalled fork after replication stress, and that AURKB does not affect double-strand breaks after restart.

### Cells require AURKB/ipl1 activity to inactivate the DDR

We determined if steps of restarting forks were inhibited by Aurora inhibition to corroborate our conclusion of a role of AURKB in restarting forks. The intra-S DDR checkpoint must be extinguished for DNA replication to restart^24–26^. The intra-S checkpoint activates the effector kinase Rad53 (yeast)/CHK1 (humans) by phosphorylation and dephosphorylation is a direct measure of extinguishing the checkpoint. We measured the phosphorylation of Rad53 in wild type, *ipl1-321* and *cdc5-2* yeast cells by mobility shift of HA-tagged Rad53 assayed by immunoblotting. Cycling cells were treated with 0.01% MMS for forty-five minutes at the restrictive temperature for *ipl1-321* and *cdc5-2* and then the MMS was removed. Approximately 50% of the Rad53 was hyperphosphorylated in wild type cells and became completely dephosphorylated three to four hours after removing the MMS (Figure 4A). In contrast, *ipl1-321* and *cdc5-2* cells retained high levels of hyperphosphorylated Rad53 suggesting that the mutants could not extinguish checkpoint signaling. We confirmed that the change in electrophoretic mobility corresponded to a change in protein kinase activity by directly measuring the protein kinase activity *in situ*^27^. We renatured Rad53 after immunoblotting and incubated the membranes with ^32^P-ATP to detect autophosphorylation of Rad53. The activity of the Rad53 kinase matched the phosphorylation state and wild type cells activated and then deactivated the kinase but the *ipl1-321* and *cdc5-2* cells retained active kinase (Supplemental Figure 4A). HeLa cells in prolonged replication stress had increased phosphorylation of CHK1 at two distinct ATM/ATR sites, S345 and S317 (Figure 4B, Supplemental Figure 4C). Phosphorylation of S345 declined over time after release from replication stress and returned to baseline after two hours. Inhibiting either AURKB or PLK1 resulted in increased and prolonged phosphorylation of CHK1 S345 and baseline signaling did not return after 4 hours (Figure 4B right panel). We obtained a similar result for the phosphorylation of CHK1 S317 (Supplemental Figure 4B and C). AURKB and PLK1 promote recovery from replication stress in an evolutionarily conserved manner.

**Figure 4.**
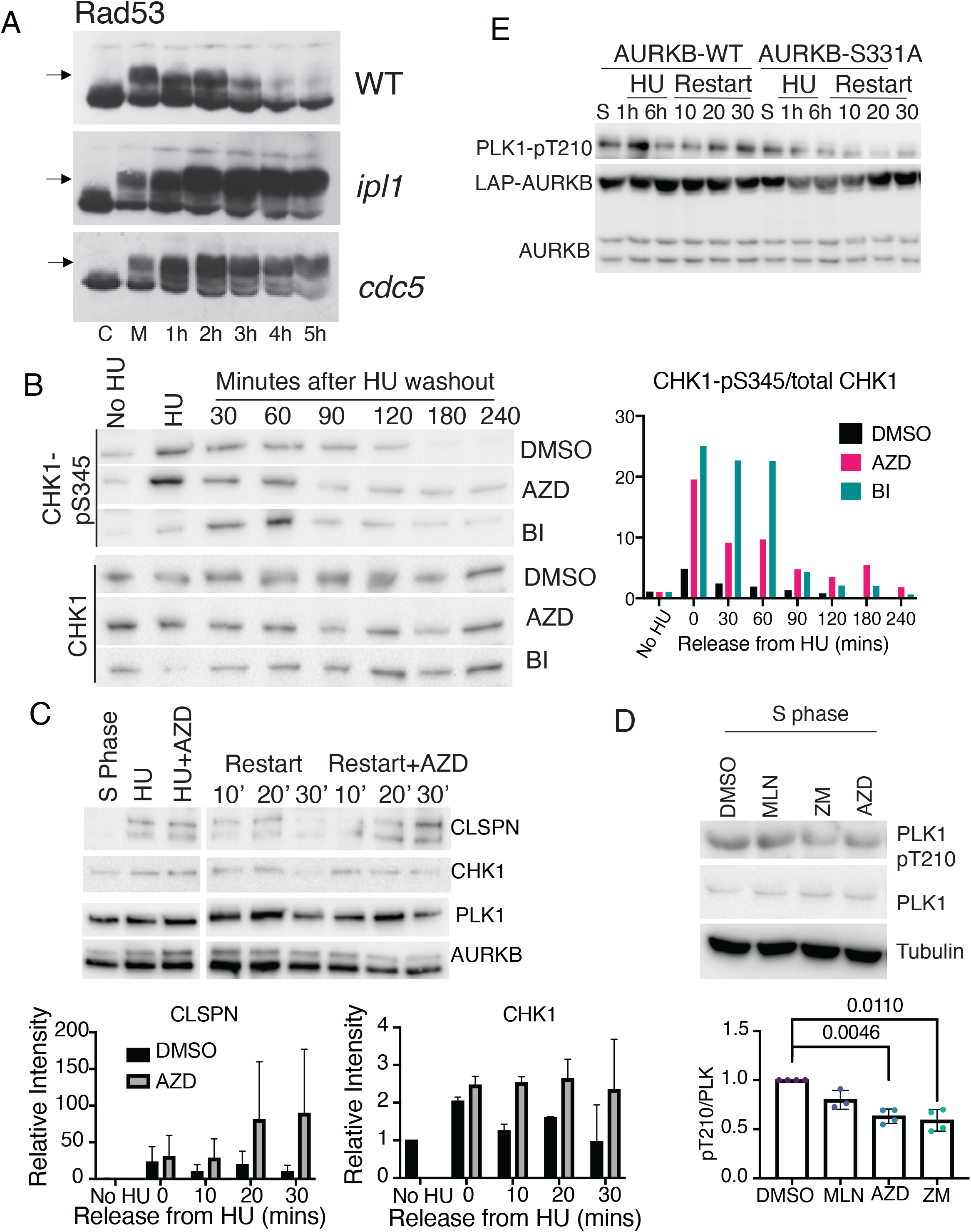
Aurora B acts through PLK1 to turn off the DNA damage checkpoint signal and release the fork protection complex from chromatin. Replication restart requires CHK1 activation of AURKB to activate PLK1 to inactivate CHK1. **(A-B)** AURKB/Ipl1 and PLK1/Cdc5 regulates ATR/Mec1 dephosphorylation. **A**. Cycling (labeled C) wild type (4159), *ipl1-321* (4166-1-4) and *cdc5-2* (4189-14-3) cells tagged with 3xHA were treated with MMS for 45 min at 36 degrees (labeled M), the MMS was removed and cells were grown in YPD for the indicated times (hours). Proteins were separated by PAGE and HA was detected by immunoblotting. The arrow indicates hyperphosphorylated Mec1. **B**. HeLa cells were treated with HU overnight (HU, 3 mM) and washed into medium lacking HU (HU washout) containing either DMSO vehicle as a control or the AURKB inhibitor AZD-1152, 500 nM, or the PLK1 inhibitor BI-2536, 100 nM, for the indicated time and cells processed by immunoblot to visualized CHK1 phosphorylated on S345 or total CHK1. Right panel, CHK1-pS345/total CHK1, normalized to no HU control. **C**. Cells require AURKB kinase to remove Claspin (CLSPN) from chromatin during restart. Cells were synchronized into S phase, treated with 6 hours of HU with and without AZD (500 nM), and washed out of HU to restart forks in the presence or absence of AZD at the same concentration. Cells were then fractionated to yield the chromatin bound fraction and immunoblotted visualize the amount of the various proteins bound to chromatin. Quantification for Claspin (CLSPN) and CHK1 normalized to No HU control, lower panels. **D**. AURKB phosphorylates PLK1 T-loop during S phase. S-phase synchronized cells were treated with the AURKB inhibitors AZD at 500 nM or ZM at 2 μM or AURKA inhibitor MLN at 100 nM and active PLK1 was visualized by immunoblot to detect phosphorylated T-loop of PLK1. Quantification for PLK1-pT210/PLK1, normalized to DMSO control. **E**. HeLa cells conditionally expressing either LAP-AURKB (WT) or LAP-AURKB (S331A) were treated as indicated and the amount of active PLK1 assayed by immunoblot.

Part of the intra-S checkpoint is the removal of Fork Protection Complex proteins such as Claspin, Timeless, Tipin from the fork after damage is resolved. Both Claspin and CHK1 remain on chromatin upon replication restart in the presence of AURKB inhibitors as opposed to AURKB, PLK1, and BUB1 that remain on chromatin in both conditions (Figure 4C, PLK1 quantified in Supplemental Figure 4D).

Aurora B phosphorylates Histone H3 on Serine 10 during replication restart (Figure 1). To corroborate a role of AURKB in restart we searched for additional substrates that have been implicated in silencing the DDR. AURKA kinase abrogates the DDR just before cells enter mitosis by activating PLK1 through direct phosphorylation of its T-loop on T210^28,29^. We determined whether AURKB activates PLK1 during the restart of replication forks in S phase. Inhibiting AURKB, but not AURKA, reduced PLK1 activation as measured by immunoblot in S phase (Figure 4D). PLK1-pT210 was also increased in HU and during replication restart (Figure 4E, Supplemental Figure 4C). The high activity in restart was abrogated by AURKB inhibition (Supplemental Figure 4C). CHK1 phosphorylates AURKB on S331 to fully activate its activity^13^. We generated cells expressing either WT or S331A AURKB behind an inducible promoter. Expression of AURKB S331A, but not wild-type AURKB, reduced PLK1-T210 phosphorylation in HU and during restart (Figure 4E). AURKB/Ipl1 activates PLK1/Cdc5 to extinguish the DDR and recover from replication stress.

## Discussion

We have uncovered an unappreciated, highly conserved function of the CPC in recovery from replication stress. Replication forks fail to restart in the absence of AURKB activity and the DDR is not turned off after the stress is removed. We have provided evidence for an extensive signaling mechanism involving known CPC regulators at stalled replication forks. The regulators that recruit (BUB1 and HASPIN) and activate (CHK1) at inner centromeres are also required to recruit and activate CPC at stalled forks. AURKB is active at forks since it is phosphorylating Histone H3 on S10 and PLK1 on its activation loop. We have provided evidence that PLK1/Cdc5 is also required to inactivate the DDR in yeast and essentially phenocopies Ipl1 mutant in their inability to inactivate Rad53 kinase after recovering from replication stress. While demonstrating that each of these steps is required for restarting forks is beyond the scope of this short report, we suggest that elucidating this pathway is an important area of future research.

AURKB kinase inhibitors are currently being tested as cancer therapeutics with variable responses^30^. Interestingly, there was over 50% positive responses when the AURKB inhibitor Barasertib (AZD1152) was used in combination with low dose cytosine arabinoside in elderly patients with AML ^31^. Our data provide an important re-interpretation of this result as we can increase cytotoxicity of AURKB kinase inhibitors during S-phase in cells that never reach mitosis while the kinase is inhibited. A major limitation of AURKB kinase inhibitors as chemotherapeutics has been the side effect neutropenia at the doses of inhibitors required to elicit mitotic defects. Thus, we suggest that re-purposing AURKB inhibitors in combination with DNA damaging agents will allow use of these inhibitors at lower doses and enable a productive line of future cancer therapeutics.

## Methods

### Yeast Strains and Media

Rad53 was epitope-tagged with three tandem hemagglutinin (3HA) sequences using a marker swapping plasmid pCMY87 provided by Dr. Chris Yellman (University of Texas). A DNA fragment (3HA kanmx) was PCR amplified using *RAD53* C-terminal tagging primers. Gel purified DNA was transformed into TAP-tagged Rad53 (Sp his5^+^) cells from the genome TAP collection and transformants were selected that were G418 resistant and his^-^. *RAD53-3HA* was amplified from the strain using the flanking primers GGACCAAACCTCAAAAGGCCCCG and GAATTCTGAGTATTGGTATCTACCATCTTCTCTC followed by transformation into wild type (CVY43) and *ipl1-321* (2948-3-3) cells. The presence of the 3HA tag was confirmed by PCR using flanking primers GATCCTAGTAAGAAGGTTAAAAGGGC, CAAAACGTCACTCTATATGTAATAAAAACCC and by immunoblotting protein extracts from cycling cells with 12CA5 monoclonal anti-HA antibody.

The lithium acetate method was used for transformation and transformants were selected on YPD+G418 plates^32^. The ATP analogue 1-naphthyl-pyrazolo[3,4-d]pyrimidine (NAPP, Cayman Chemicals) was added to YPD at 75 μM. Hydroxyurea and methyl methanesulfonate (Sigma) were supplemented in the media as noted in the figure legends. Spotting assays were performed by growing cells to saturation and spotting serial ten-fold dilutions onto plates and incubating the plates at the indicated temperatures with plates sealed and submerged in waterbaths. Spotted cells on solid medium were irradiated with UV light in a Stratalinker (Stratgene), wrapped in aluminum foil and incubated at the indicated temperatures.

### Yeast Cell Cycle Experiments

Cells were grown overnight in YPD to mid-log, adjusted to OD=1.0 and synchronized at 23 degrees in YPD medium containing 100 nM of α-factor (Genway Biotech Inc) for three hours. Efficiency of the arrest was monitored by microscopy to obtain >85% as unbudded cells. Cultures of temperature sensitive mutants were incubated for the last 30 minutes at 36 degrees to inactivate the protein. Arrested cells were washed three times with water and released into YPD medium in the presence or absence of MMS at temperatures indicated in the figure legends. Samples were collected at the indicated times, sample were fixed with 70% ethanol and prepared for flow cytometry as described previously^33^. Flow cytometry was performed with a BD FACScalibur flow cytometer in the core facility at the University of Virginia or with a BD Accuri C6 flow cytometer at North Carolina State University. The DNA content of 40,000 cells was determined for each sample. Rad53 hyperphosphorylation was detected by immunoblotting as previously described^33^. Cells were grown to mid-log at 23 degrees in YPD and adjusted to OD=1. MMS (0.01%) was added for 45 minutes and quenched with 0.5% sodium thiosulfate. Cells were pelleted, washed with water and resuspended in pre-warmed YPD and grown at 36 degrees. Protein extracts were prepared as described previously and immunoblotted with mouse monoclonal 12CA5 antibody. Real Time PCR was performed with a StepOne system (Applied Biosystems) following manufacturer’s instructions. Primers to amplify early ARS sequences: ARS1 (TTTCTGACTGGGTTGGAAGG, CGCATCACCAACATTTTCTG), ARS305 (GATTGAGGCCACAGCAAGAC, TCACACCGGACAGTACATGA) and late sub-telomeric ARS sequences: ARS440 (CGAAAGTGACGAAGTTCATGC, GCCATTGCTGATAAAGACGC), ARS522 (GTTTTAGCAGCTCCAAAAGAAAGG, GGACTTTAGATAGTAATATATGGCG). Standard curves were prepared for each primer pair using purified genomic DNA quantified using nanodrop and absolute amounts of each amplified PCR product was determined from the standard curves.

*In situ* protein kinase assay on lysates from wild type and *ipl-321* cells was performed as outlined in^27^. Yeast proteins were loaded onto an 8% electrophoresis gel and blotted onto a PVDF membrane, denatured at room temperature for 1 hour and renatured overnight at 4C. The membrane was washed in 30mM Tris pH 7.5 for one hour and incubated in kinase buffer for 30min. 10 μCi of γ-32 P was added to fresh kinase buffer and incubated at room temperature for one hour. The blot was washed in 30mM Tris, followed by 1M KOH for 10min and 10%TCA for 10min. Activity was detected via autoradiography.

### Human Cell Culture and Drug Treatments

Cells were synchronized into S phase by double thymidine block and release: 2mM thymidine (Sigma-Aldrich) was added to culture medium for 24 hours, washed out into normal medium for 12 hours, then treated with 2mM thymidine-containing medium for 12 hours. Cells were released to into medium for 1-2 hours to resume S-phase before the addition of drugs. ZM44739 (Enzo Life Sciences) was dissolved in DMSO and cells were treated with 500nM. AZD1152 (MedChemExpress) were used at concentrations of 100nM. Control cells were treated with identical amounts of DMSO carrier. HeLa cells were checked for mycoplasma by PCR biannually. Immunofluorescence was performed as described (Bolton et al. 2002).

### Colony Assay

HeLa T-Rex cells were synchronized into S Phase and treated with 2mM Hydroxyurea and DMSO or Aurora Inhibitors for 4, 6, or 8 hours before being rinsed in PBS and put in DMEM overnight at 37°C. The following day, cells were trypsinized and counted with a coulter counter and diluted and 50 cells were transferred to a new plate and cultured for 12-14 days when colonies were visible in untreated samples. The plates were fixed in 100% methanol for 20min at room temperature and stained with Crystal Violet (0.5% in 25% methanol). Colonies were counted manually.

### Human Cell Line Generation

The S331A cells were generated through Quick Change site Directed Mutagenesis of previously made PcDNA5.0/FRT/TO-LAP-hAuroraB^34^. Tet-inducible cell lines were generated as described^34^.

### Immunoblotting

Immunoblotting was performed as described^35^. Images were collected through autoradiography or imaged on a BioRad ChemiDoc. The antibodies used were: 12Ca5 (mouse monoclonal antiHA produced in UVA hybridoma facility), Borealin (Banerjee JCB 2014), AURKA (homemade), H3S10P (Mouse Millipore Cat#06570 multiple Lot#, Rabbit Cell Signaling Technologies Cat#D2C8 Lot# 3377S), H3T3 (EMD Millipore Cat#7-424 Lot#3012075), H2AP.T120 (Active Motif Cat#61195 Lot#31511001), BUB1 (GeneTex Cat#GTx30097 Lot#821701160, PCNA (Novus Biotechne-Brand NB500-106 Lot # A6), H2B (Upstate Cat#07371 Lot#30216) H4 (Cell Signaling Technologies), PLK1P.T210 (Cell Signaling Technologies Lot#A3730), Tubulin (DM1a, UVA hybridoma facility), P-CHK1-S345(Cell Signaling Cat#133D3, Lot#2348S), P-CHK1-S317 (Cell Signaling Cat#2344S) P-ATR90, antiBrdU(Mouse-BD Biosciences 347580 Lot# 7157935 AbCam Rat Cat#AB6326 multiple lots #GR3173537-7, #GR3269246-1), gamma-H2aX (Upstate Cell Signaling Solutions Cat# 07-164 Lot#23646), Pan Aurora T-loop (Cell Signaling Technologies), Claspin (A300267A-Bethyl), IdU (from Burke), CldU (from Burke). All secondary antibodies were from Jackson Labs. Aurora A/B/C-P (Cell Signaling Technologies-Cat#2914S Lot#9) Timeless (Sigma SAB1406756-50ug Lot# 08242), PLK1 (Cell Signaling Technologies Cat#4513S), Total CHK1 (Santa Cruz, SC8408 Lot#K0118), Rabbit AURKB Total (Bethyl-A300431A).

### aniPOND

HeLa TRex cells were grown to 90% confluency at 37□C and 5% CO2 to obtain at least 1×10^8^ cells per treatment to perform aniPOND as described in^22^. Briefly, cells were treated with 10 μM EdU (LifeTechnologies) for ten minutes or with 2mM HU with 10 μM EdU for six hours in a 37°C incubator. EdU containing media was replaced in chase samples with medium containing 10 μM thymidine and incubated for twenty minutes in 37°C. The HU containing medium in restart experiments was washed out and replaced with new medium for twenty minutes with fresh EdU (and AZD/ZM where mentioned) at 37°C. Nuclei were extracted and isolated on ice. The EdU was conjugated to biotin through a Click Reaction. When cells number per sample exceeded 1×10^8^, 10mLs of click reaction buffer was used, otherwise 5mL of solution was used, with final concentrations 25 μM biotin-azide, 10 mM (+)-sodium L-ascorbate, 2 mM CuSO4 and rotated at 4°C for 1-2hr. Samples were then sonicated on ice using a Branson Sonifier-250 for 12 pulses each 10 seconds with 20W. Some of the sample was set aside as the “input” sample. The rest of the sample was incubated with streptavidin-coated agarose beads overnight at 4°C. Beads were spun down and washed before the proteins were eluted off the beads by boiling in SDS sample buffer and processed for immunoblot.

### Cell Fractionation

Chromatin-bound samples were isolated using a modified version of the cell fractionation protocol outlined in^36^. Cells were lysed in Buffer A (10 mM HEPES pH 7.9, 10 mM KCl, 1.5 mM MgCl2, 34 mM sucrose, 10% glycerol, 1 mM DTT, 5 mM Na3VO4, 10 mM NaF, protease inhibitor tablet) with 0.1% Triton X-100. The supernatant was isolated after a 4min spin at 1300g and labeled as the “cytoplasmic fraction”. The pellet was treated with Buffer B (3 mM EDTA, 0.2 mM EGTA, 1 mM DTT, 5 mM Na3VO4, 10 mM NaF, protease inhibitor tablet) for 1 minute on ice and spun at 1700x g for 4 minutes. The resulting pellet was resuspended in Buffer A with 1mM CaCl2 and 5 U micrococcal nuclease and incubated at 37°C for 10 minutes. The chromatin bound fraction was collected as the supernatant after a spin at max speed for 5 minutes.

### DNA Combing and Fiber Spreading assays

DNA fiber spreading was performed as described^37^. Cells were pulsed with 20 μM IdU for twenty minutes, then the media was removed, the cells were washed twice with Hanks balanced salt solution (HBSS, Fisher), then treated with HU for 6 hours and 100 μM CldU for twenty minutes. The cells were then trypsinized, spun at 800 x g and washed in PBS and resuspended in a volume to bring the final concentration to approximately 1 × 10^6^ cells/ml. 2 μl of the suspension was place on a clean slide and lysed with 10 μl of lysis buffer (0.5% SDS, 200mM Tris pH 7.4, and 50mM EDTA) for 6 minutes. Then the slides were tilted to allow the solution to drip slowly down the length of the slide, and then air dried at room temperature. Slides were fixed in a 3:1 methanol: acetic acid solution for two minutes, then air dried again. The slides were stored overnight a 4°C and the DNA was denatured in 2.5 M HCl for 30 minutes, then rinsed 3 times in PBS. The slides were incubated with blocking buffer (BB, 10% Goat serum, 0.1% Triton, PBS) for 1 hour and incubated in rat anti-BrdU diluted 1:300 in BB. Subsequently, the slides were incubated for an hour in a 1:100 dilution of mouse anti-BrdU. Alexa-Fluor antibodies were used at a dilution of 1:350. Slides were air dried then mounted with ProLong Gold.

## Data Availability

Data supporting the findings of this work are available within the paper and its Supplementary Information files. The datasets are available from the corresponding author upon request.

## Supplementary Figure Legends

**Supplemental Figure 1.**
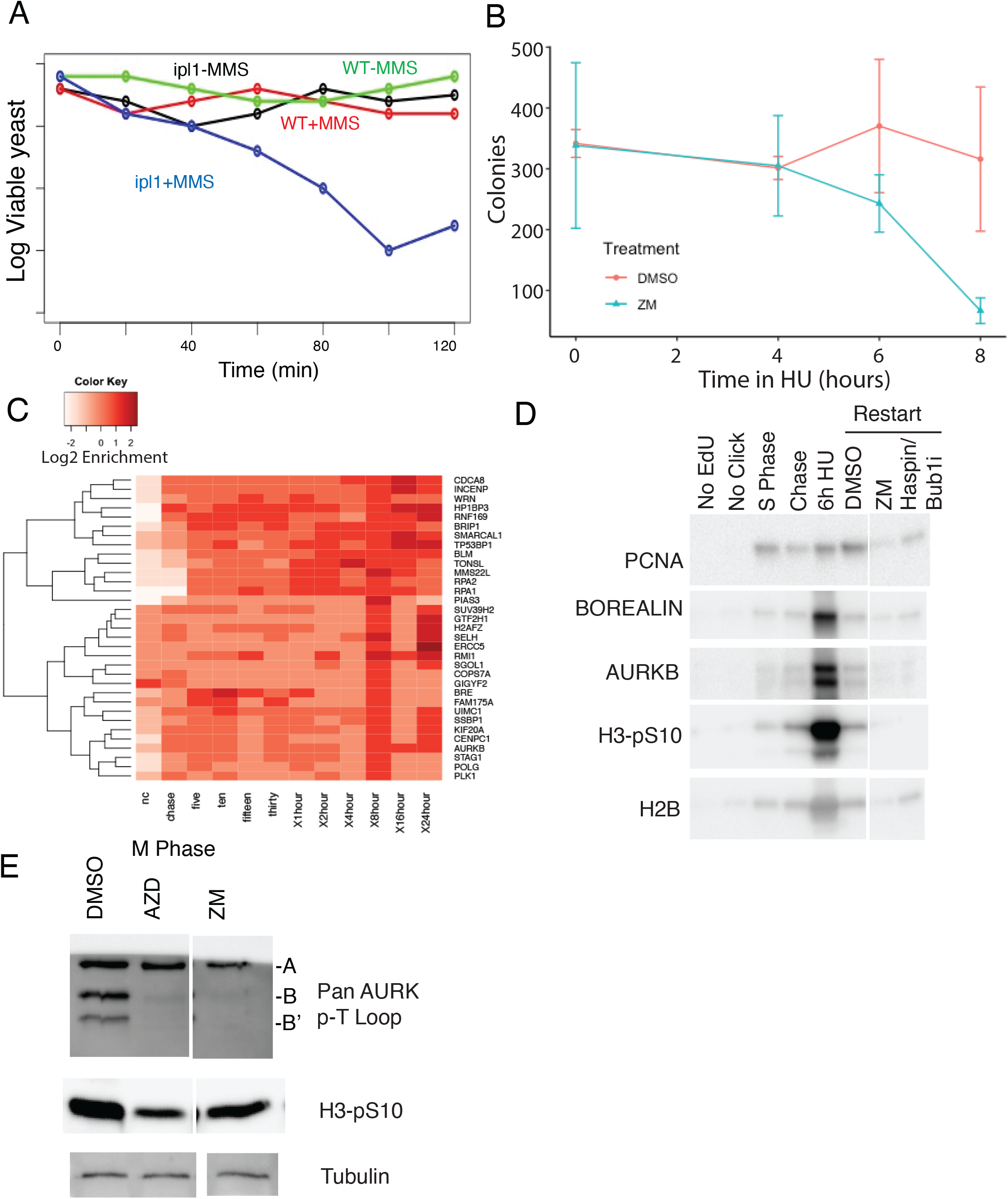
AURKB has a role in the response to DNA damage. **A**. Plating efficiency. Wild type (CVY43) and *ipl1* (2948-3-3) cells (10^7^ per ml) were treated with 0.033% MMS for the indicated time, diluted onto YPD plates and counted to determine the number of viable cells. **B**. HeLa cells fail to form colonies after being treated with HU and Aurora inhibitors. Cells were treated for the indicated time and drugs diluted and replated at the same cell number and then grown for 14 days in fresh media to form colonies. p-value=0.04 (Anova). **C**. Hierarchical clustering of quantitative iPond data. Clustering of Stable Isotope Labeling with Amino acids in Cell culture (SILAC) data for proteins that accumulate late at replication forks. **D**. The CPC is preferentially bound to replication-stressed chromatin in HeLa cells synchronized in S-phase as measured by iPOND. Cells synchronized by thymidine were treated with EdU for 10 minutes (S-phase), or chased with 10 uM thymidine for 20 minutes, newly replicated chromatin was isolated on streptavidin beads after treatment of chromatin with click-it biotin and the isolated chromatin was analyzed by immunoblot with the indicated antibodies. **E**. The AURKB inhibitors do not inhibit AURKA kinase at the concentrations used in this study. Nocodazole-arrested cells were treated with100nM AZD (AZD) or 400nM ZM447439 (ZM) for 1 hour and the amount of active AURKA (46 KD) and AURKB (41, 36KD) kinases measured by immunoblot against a Pan Aurora phospho-T-loop antibody and antibodies against an AURKB substrate Phosphoserine-10 of histone H3 (H3-pS10).

**Supplemental Figure 2.**
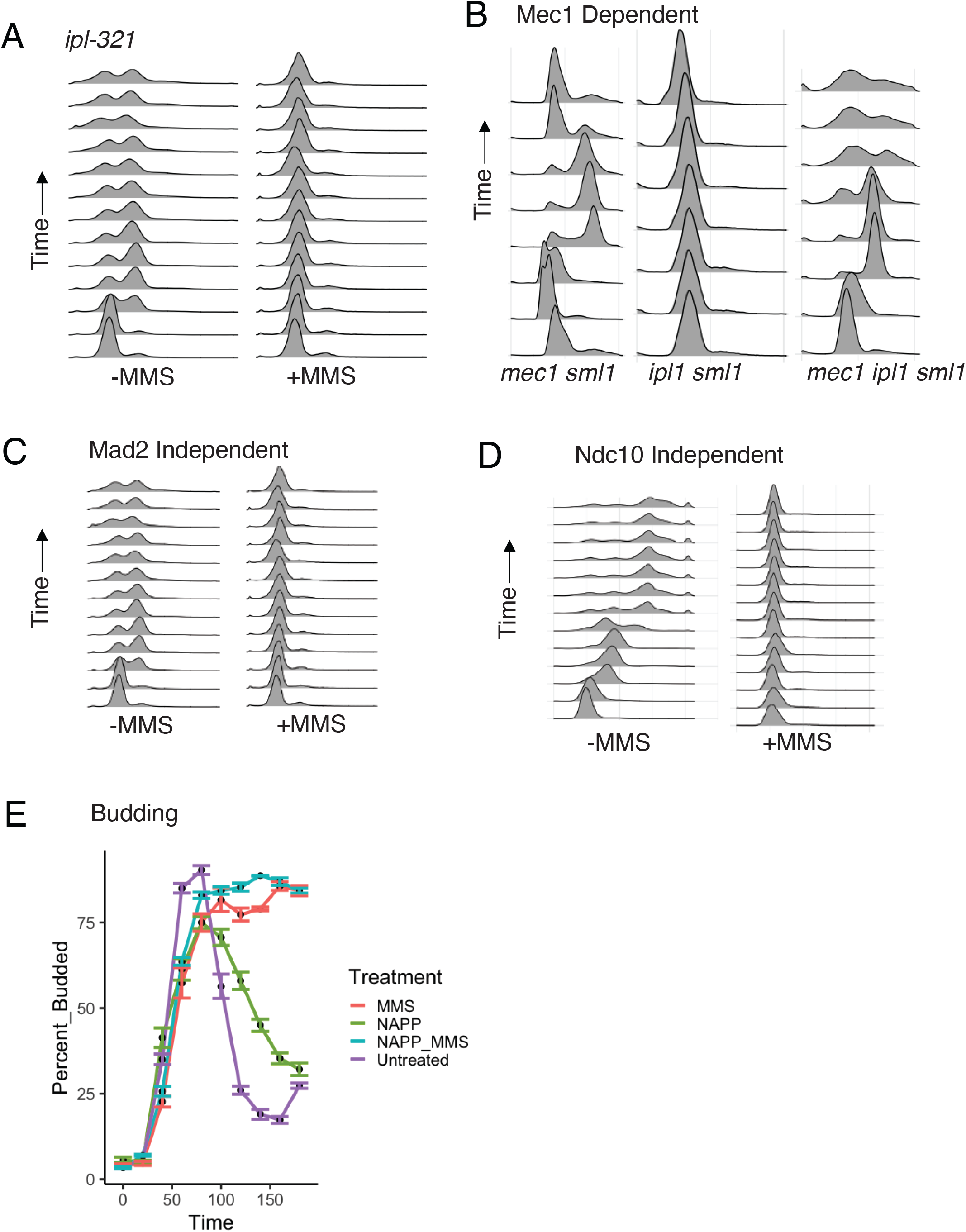
Ipl1 cells respond to MMS through the DNA damage checkpoint. **A**. *ipl-321* do not complete DNA replication in MMS. α-Factor-arrested *ipl1-321* cells (4145-1-3) were released into the cell cycle for the indicated time at 36 degrees and samples were taken every twenty minutes for flow cytometry. **B**. The Ipl1 response to replication stress requires Mec1. α-Factor-arrested *ipl1-321 sml1 (4145-1-3), mec1 sml1* (4161-10-2), and *ipl1-321 mec1 sml1* (4145-10-4) cells were released into the cell cycle in 0.033% MMS at 36 degrees for the indicated time and samples were taken every 20 min for flow cytometry. **C**. The Ipl1 response to replication stress does not require Mad2. α-Factor-arrested *mad2::kanmx4* cells (2492-6-1) were released into the cell cycle in the presence or absence of 0.033% MMS at 36 degrees for the indicated time and samples were taken every 20 min for flow cytometry. **D**. The Ipl1 response to replication stress does not require the kinetochore. α-factor-arrested *ndc10-1* cells (2495-12-3) were released into the cell cycle in the presence or absence of 0.033% MMS at 36 degrees for the indicated time and samples were taken every 20 min for flow cytometry. **E**. Budding data from experiment shown in Figure 2A to show cells had entered S-phase under all conditions.

**Supplemental Figure 3.**
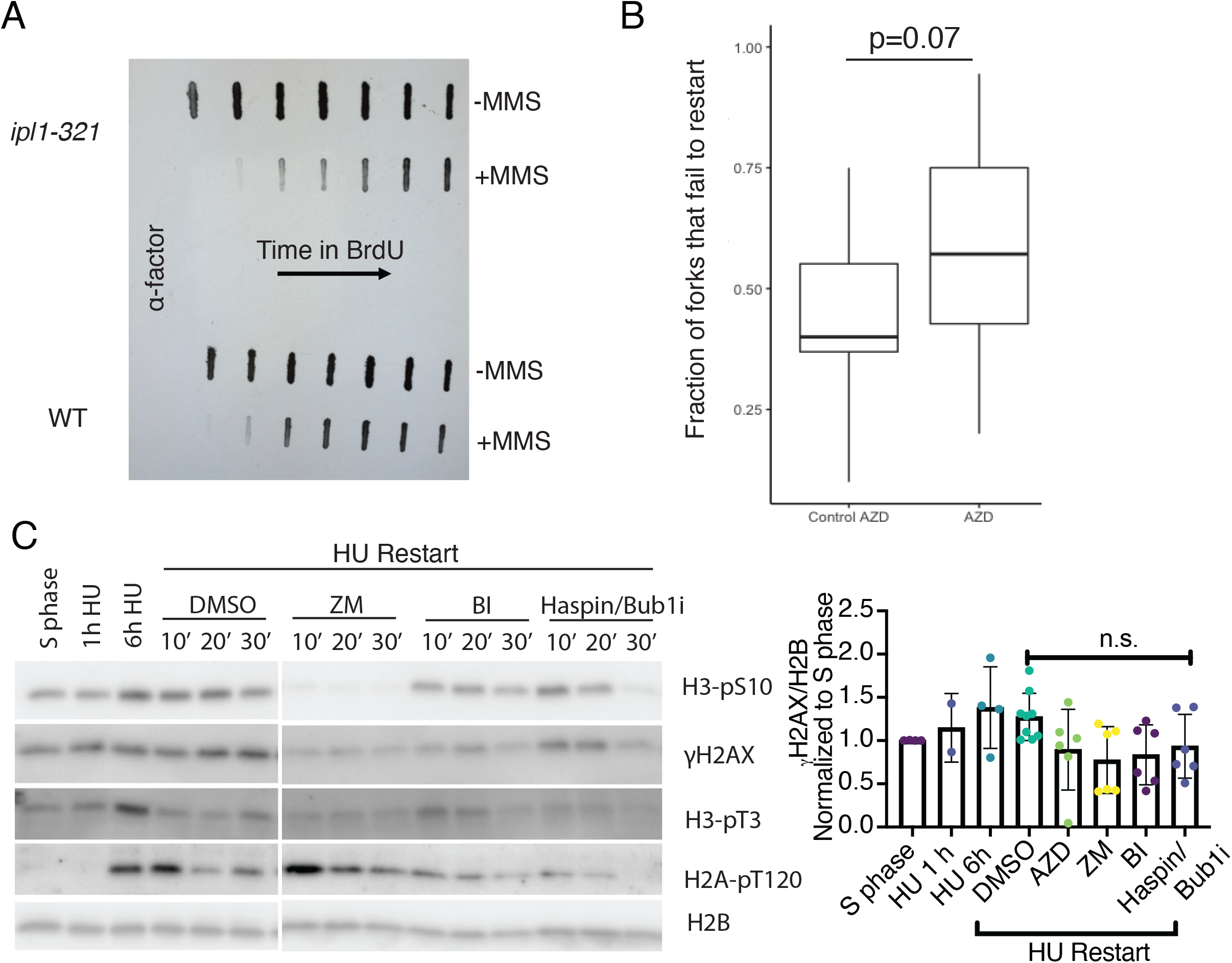
AURKB is required for efficient replication restart. **A**. Wild type (CVY43) and *ipl1-321* (2948-3-3) cells were α-factor-arrested are released into YPD plus BrdU and sampled every 20 minutes. The +MMS indicates cells treated with 0.033% MMS for 30 min after α-factor release and then the MMS was removed to monitor replication restart. Equal amounts of DNA (nanodrop) were denatured, captured by slot blot and BrdU was detected by immunoblotting. **B**. Replication from restarted replication forks was measured as in Figure 3C except cells were treated with the AZD AURKB inhibitor rather than ZM and assayed by DNA fiber spreading rather than combing. **C**. Restart experiment was performed by stalling S-phase synchronized cells in 3 mM HU for 6 hours, then washing out into the labeled inhibitor ant taking whole cell extracts at 10, 20, and 30 minutes. ZM, 2 μM, BI, 100 nM, Haspin inhibitor (5ITU), 1 μM, BUB1 inhibitor (BAY-1816032), 1 μM.

**Supplemental Figure 4.**
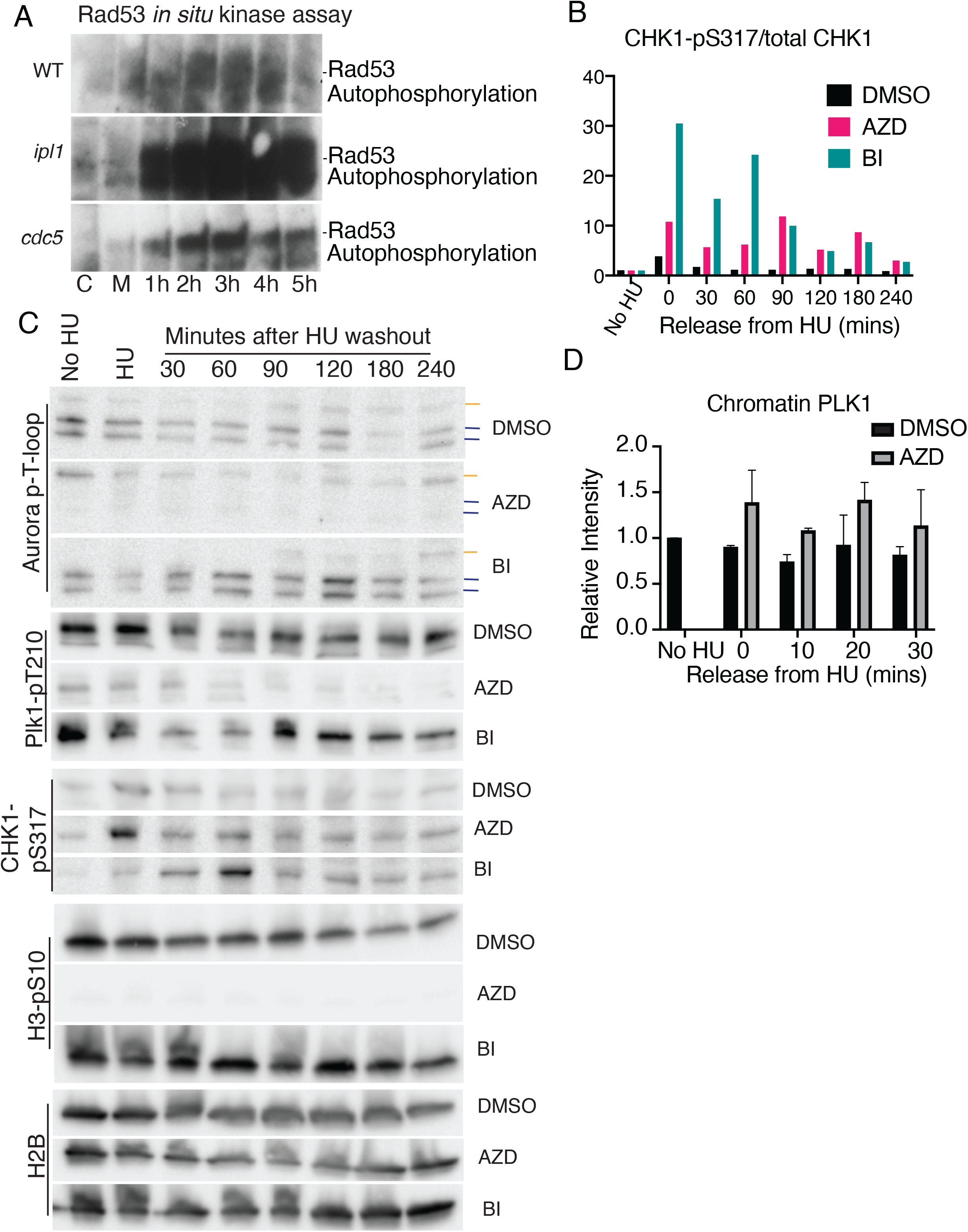
AURKB/Ipl1 and PLK1/Cdc5 regulate ATR/Mec1. **A**. *In situ* kinase assay of cycling wild type, *ipl1-321* and *cdc5-1* cells tagged with 3xHA (labeled C) treated with MMS for 45 min at 36 degrees (labeled M), the MMS was removed and cells were grown in YPD for the indicated times (hours). Proteins were separated by PAGE and Mec1 kinase activity was detected by autophosphorylation after kinase renaturation and incubating with ^32^P-labelled ATP followed by autoradiography. **B**. Quantification of PLK1 T-loop phosphorylation during S phase after Aurora kinase inhibition. **C**. Cells were treated as in Figure 4B, then immunoblotted for pan-Aurora T-loop phosphorylation (AURKA, yellow, AURKB, blue), PLK1 pT210, H3-pS10, and H2B as a loading control. **D**. Quantification of chromatin bound Claspin (CLSPN), CHK1, and PLK1. Claspin and CHK1 stay bound to chromatin across multiple experiments after AURKB inhibition, while chromatin bound PLK1 is unaffected by AURKB inhibition.

## Acknowledgements

We thank Dr. Oscar Aparicio (USC), Dr. Jasper Rine (U.C. Berkley), Dr. Sue Biggins (FHCRC), Dr. Chris Yellman (U.T.) and Dr. Yanchang Wang (FSU) for yeast strains and plasmids. We would like to also thank Dr. Etsuko Shibata and Dr. Anindya Dutta for help with the combing experiments. We especially thank Dr. David Cortez for pointing out that AURKB was at stalled replication forks and for constructive conversations throughout the work. KP and EM were supported by the Cancer Training Grant at UVA (T32CA009109) and overall, the work was supported by R01 GM118798 to PTS and DJB.

## Author Contributions

Yeast work was performed by BV under the supervision of DJB. Most work on human cells was performed by EM with initial experiments done by KP under the supervision of PTS. SKM performed intial experiments for 3C. The paper was written by EM, PTS and DJB.

**Supplemental Table 1.** Yeast Strains. All strains are isogenic with W303 (*MAT***a** *ade2-1 his3-11,15 leu2-3,112 trp1-1 ura3-1 can1-*100) and indicated genotypes are the additional mutations.

| Strain | Genotype | Source |
| --- | --- | --- |
| CVY43 | *bar1::hisG URA3:BrdU-inc* | Oscar Aparicio |
| JRY7274 | *mec1::TRP1 sml1-1* | Jasper Rine |
| 2492-6-1 | *bar1::hisG ipl1::KANMX:ipl1-as5:LEU2*<br>*mad2::KANMX* | This Study |
| 2495-12-3 | *ndc10-1 ipl1-321* | This Study |
| 2498-3-3 | *bar1::hisG URA3:BrdU-inc ipl1-321* | This Study |
| 2946-14-2 | *bar1::hisG ipl1::KANMX:ipl1-as5:LEU2 URA3:BrdU-inc* | This Study |
| 4145-1-3 | *sml1::HIS3 ipl1-321* | This Study |
| 4145-10-4 | *mec1::TRP1 sml1::HIS3 ipl1-321* | This Study |
| 4159 | *bar1::hisG URA3:BrdU-inc RAD53-3HA:KanMX* | This Study |
| 4161-10-2 | *bar1::hisG ipl1-321 mec1::TRP1 sml1* | This Study |
| 4166-1-4 | *bar1::hisG URA3:BrdU-inc ipl1-321 RAD53-3HA:KanMX* | This Study |
| 4189-14-3 | *cdc5-2 RAD53-3HA:KANMx4* | This Study |

## References

1. Técher, H., Koundrioukoff, S., Nicolas, A. & Debatisse, M. The impact of replication stress on replication dynamics and DNA damage in vertebrate cells. Nature Reviews Genetics (2017) doi:10.1038/nrg.2017.46.

2. Branzei, D. & Foiani, M. Maintaining genome stability at the replication fork. Nature Reviews Molecular Cell Biology (2010) doi:10.1038/nrm2852.

3. Cortez, D. Preventing replication fork collapse to maintain genome integrity. DNA Repair (2015) doi:10.1016/j.dnarep.2015.04.026.

4. Saldivar, J. C., Cortez, D. & Cimprich, K. A. The essential kinase ATR: Ensuring faithful duplication of a challenging genome. Nature Reviews Molecular Cell Biology (2017) doi:10.1038/nrm.2017.67.

5. Trivedi, P. & Stukenberg, P. T. T. A Centromere-Signaling Network Underlies the Coordination among Mitotic Events. Trends in Biochemical Sciences vol. 41 (2016).

6. Pfister, K. et al. Identification of Drivers of Aneuploidy in Breast Tumors. Cell Rep. 23, (2018).

7. Tanno, Y. et al. The inner centromere-shugoshin network prevents Chromosomal instability. Science (2015) doi:10.1126/science.aaa2655.

8. Carmena, M., Wheelock, M., Funabiki, H. & Earnshaw, W. C. The chromosomal passenger complex (CPC): From easy rider to the godfather of mitosis. Nat. Rev. Mol. Cell Biol. (2012) doi:10.1038/nrm3474.

9. Wang, F. et al. Histone H3 Thr-3 phosphorylation by haspin positions Aurora B at centromeres in mitosis. Science 330, (2010).

10. Kelly, A. E. et al. Survivin reads phosphorylated histone H3 threonine 3 to activate the mitotic kinase Aurora B. Science (2010) doi:10.1126/science.1189505.

11. Liu, H., Jia, L. & Yu, H. Phospho-H2A and cohesin specify distinct tension-regulated sgo1 pools at kinetochores and inner centromeres. Curr. Biol. (2013) doi:10.1016/j.cub.2013.07.078.

12. Kawashima, S. A., Yamagishi, Y., Honda, T., Lshiguro, K. I. & Watanabe, Y. Phosphorylation of H2A by Bub1 prevents chromosomal instability through localizing shugoshin. Science (2010) doi:10.1126/science.1180189.

13. Petsalaki, E., Akoumianaki, T., Black, E. J., Gillespie, D. A. F. & Zachos, G. Phosphorylation at serine 331 is required for Aurora B activation. J. Cell Biol. (2011) doi:10.1083/jcb.201104023.

14. Macůrek, L. et al. Polo-like kinase-1 is activated by aurora A to promote checkpoint recovery. Nature (2008) doi:10.1038/nature07185.

15. Pellicioli, A., Lee, S. E., Lucca, C., Foiani, M. & Haber, J. E. Regulation of Saccharomyces Rad53 checkpoint kinase during adaptation from DNA damage-induced G2/M arrest. Mol. Cell (2001) doi:10.1016/S1097-2765(01)00177-0.

16. Toczyski, D. P., Galgoczy, D. J. & Hartwell, L. H. CDC5 and CKII control adaptation to the yeast DNA damage checkpoint. Cell (1997) doi:10.1016/S0092-8674(00)80375-X.

17. Yoo, H. Y., Kumagai, A., Shevchenko, A., Shevchenko, A. & Dunphy, W. G. Adaptation of a DNA replication checkpoint response depends upon inactivation of Claspin by the Polo-like kinase. Cell (2004) doi:10.1016/S0092-8674(04)00417-9.

18. Wakida, T. et al. The CDK-PLK1 axis targets the DNA damage checkpoint sensor protein RAD9 to promote cell proliferation and tolerance to genotoxic stress. eLife (2017) doi:10.7554/eLife.29953.

19. Yata, K. et al. Plk1 and CK2 Act in Concert to Regulate Rad51 during DNA Double Strand Break Repair. Mol. Cell (2012) doi:10.1016/j.molcel.2011.12.028.

20. Li, Z. et al. Plk1 phosphorylation of Mre11 antagonizes the DNA damage response. Cancer Res. (2017) doi:10.1158/0008-5472.CAN-16-2787.

21. Dungrawala, H. et al. The Replication Checkpoint Prevents Two Types of Fork Collapse without Regulating Replisome Stability. Mol. Cell (2015) doi:10.1016/j.molcel.2015.07.030.

22. Thomas Leung, K. H., El Hassan, M. A. & Bremner, R. A rapid and efficient method to purify proteins at replication forks under native conditions. BioTechniques (2013) doi:10.2144/000114089.

23. Pinsky, B. A., Kotwaliwale, C. V, Tatsutani, S. Y., Breed, C. A. & Biggins, S. Glc7/protein phosphatase 1 regulatory subunits can oppose the Ipl1/aurora protein kinase by redistributing Glc7. Mol Cell Biol 26, 2648–2660 (2006).

24. Szyjka, S. J. et al. Rad53 regulates replication fork restart after DNA damage in Saccharomyces cerevisiae. Genes Dev. (2008) doi:10.1101/gad.1660408.

25. Travesa, A., Duch, A. & Quintana, D. G. Distinct phosphatases mediate the deactivation of the DNA damage checkpoint kinase Rad53. J. Biol. Chem. (2008) doi:10.1074/jbc.M801402200.

26. Hustedt, N. et al. Yeast PP4 interacts with ATR homolog Ddc2-Mec1 and regulates checkpoint signaling. Mol. Cell (2015) doi:10.1016/j.molcel.2014.11.016.

27. Pellicioli, A. et al. Activation of Rad53 kinase in response to DNA damage and its effect in modulating phosphorylation of the lagging strand DNA polymerase. EMBO J. (1999) doi:10.1093/emboj/18.22.6561.

28. Macurek, L. et al. Polo-like kinase-1 is activated by aurora A to promote checkpoint recovery. Nature 455, 119–123 (2008).

29. Seki, A., Coppinger, J. A., Jang, C.-Y., Yates, J. R. & Fang, G. Bora and the kinase Aurora a cooperatively activate the kinase Plk1 and control mitotic entry. Science 320, 1655–1658 (2008).

30. Borisa, A. C. & Bhatt, H. G. A comprehensive review on Aurora kinase: Small molecule inhibitors and clinical trial studies. European Journal of Medicinal Chemistry (2017) doi:10.1016/j.ejmech.2017.08.045.

31. H.M., K. et al. Phase i study assessing the safety and tolerability of barasertib (azd1152) with low-dose cytosine arabinoside in elderly patients with AML. Clinical Lymphoma, Myeloma and Leukemia (2013).

32. Eckert-Boulet, N., Pedersen, M. L., Krogh, B. O. & Lisby, M. Optimization of ordered plasmid assembly by gap repair in Saccharomyces cerevisiae. Yeast (2012) doi:10.1002/yea.2912.

33. Yellman, C. M. & Burke, D. J. The role of Cdc55 in the spindle checkpoint is through regulation of mitotic exit in Saccharomyces cerevisiae. Mol. Biol. Cell (2006) doi:10.1091/mbc.E05-04-0336.

34. Matson, D. R., Demirel, P. B., Stukenberg, P. T. & Burke, D. J. A conserved role for COMA/CENP-H/I/N kinetochore proteins in the spindle checkpoint. Genes Dev. 26, 542–547 (2012).

35. Bolton, M. A. et al. Aurora B kinase exists in a complex with survivin and INCENP and its kinase activity is stimulated by survivin binding and phosphorylation. Mol. Biol. Cell 13, (2002).

36. Sato, K. et al. A DNA-damage selective role for BRCA1 E3 ligase in claspin ubiquitylation, CHK1 activation, and DNA repair. Curr. Biol. (2012) doi:10.1016/j.cub.2012.07.034.

37. Bétous, R. et al. Substrate-Selective Repair and Restart of Replication Forks by DNA Translocases. Cell Rep. (2013) doi:10.1016/j.celrep.2013.05.002.

